# Range-wide habitat use and Key Biodiversity Area coverage for a lowland tropical forest raptor across an increasingly deforested landscape

**DOI:** 10.1101/2021.08.18.456651

**Authors:** Luke J. Sutton, David L. Anderson, Miguel Franco, Christopher J.W. McClure, Everton B.P. Miranda, F. Hernán Vargas, José de J. Vargas González, Robert Puschendorf

## Abstract

Quantifying habitat use is important for understanding how animals meet their requirements for survival and provides useful information for conservation planning. Currently, assessments of range-wide habitat use that delimit species distributions are incomplete for many taxa. The harpy eagle (*Harpia harpyja*) is a raptor of conservation concern, widely distributed across Neotropical lowland forests, that currently faces threats from increasing habitat loss and fragmentation. Here, we use a logistic regression modelling framework to identify habitat resource selection and predict habitat suitability based on a new method developed from the International Union for the Conservation of Nature Area of Habitat range metric. From the habitat use model, we performed a gap analysis to identify areas of high habitat suitability in regions with limited coverage in the Key Biodiversity Area (KBA) network. Range-wide habitat use indicated that harpy eagles prefer areas of 70-75 % evergreen forest cover, low elevation, and high vegetation heterogeneity. Conversely, harpy eagles avoid areas of >10 % cultivated landcover and mosaic forest, and topographically complex areas. Our habitat use model identified a large continuous area across the pan-Amazonia region, and a habitat corridor from the Chocó-Darién ecoregion of Colombia running north along the Caribbean coast of Central America. Little habitat was predicted across the Atlantic Forest biome, which is now severely degraded. The current KBA network covered ∼18 % of medium to high suitability harpy eagle habitat exceeding the target representation (10 %). Four major areas of high suitability habitat lacking coverage in the KBA network were identified in the Chocó-Darién ecoregion of Colombia, western Guyana, and north-west Brazil. We recommend these multiple gaps of habitat as new KBAs for strengthening the current KBA network. Modelled area of habitat estimates as described here are a useful tool for large-scale conservation planning and can be readily applied to many taxa.

## Introduction

Determining habitat resource use is a fundamental aspect of wildlife ecology and conservation planning (Manly *et al*. 2002; Morrison *et al*. 2006). However, our understanding of range-wide species-habitat associations across continental extents is incomplete, even for well-studied groups such as birds (Gregory & Baillie 1998; Engler *et al*. 2017; Lees *et al*. 2020). Currently, many taxa face increasing threats from human-driven habitat loss and fragmentation across their entire range (Powers & Jetz 2019). Therefore, developing a broad spatial quantification of habitat use is an effective starting point for conservation planning (Margules & Pressey 2000; Early *et al*. 2008). Once habitat use is identified for a focal species, the key variables characterising those habitats can be used to produce a mapped representation of habitat across the species’ range (Hirzel *et al*. 2006). Management actions can then be directed to guide conservation planning to protect or enhance those areas (Margules & Pressey 2000; Suárez-Seoane *et al*. 2002).

Recently, the International Union for the Conservation of Nature (IUCN) developed a new range size metric termed Area of Habitat (AOH, Brooks *et al*. 2019). AOH is defined as the habitat available to a species based on habitat preferences and elevational limits within the mapped distributional range of a focal species. Various approaches have been taken to estimate AOH which all use a similar method of matching and overlaying the known mapped range, landcover and elevation limits of a given species (Brooks *et al*. 2019). Whilst the AOH method is useful and repeatable, IUCN methods may still have limitations by missing areas that have no occurrence data but may still contain preferred habitat (Ramesh *et al*. 2017). On the other hand, Habitat Suitability Models (HSMs), and related Resource Selection Functions (RSFs), are statistical methods that assess species’ habitat requirements and predict distribution based on correlating environmental covariates with species occurrences (Boyce & McDonald 1999; Franklin 2009). Two example applications for HSMs are the re-evaluation of range sizes (e.g., Herkt *et al*. 2017), and the identification of gaps in protected or biodiversity area networks (e.g., de Carvalho *et al*. 2017). Indeed, HSMs can predict more complex and ecologically realistic geographic ranges compared to IUCN range maps (Breiner *et al*. 2017; Herkt *et al*. 2017). Using model-based interpolation based on the AOH guidelines but adapted to a correlative modelling approach (Da Silva *et al*. 2020), may also be more effective for highlighting species-specific biodiversity area gaps by identifying higher coverage of suitable pixels (Di Marco *et al*. 2017).

Biodiversity areas are a fundamental tool for conservation (IUCN 2016) and have been successful in reducing habitat loss and fragmentation for many taxa (Brooks *et al*. 2009). However, despite wide coverage in the global biodiversity area network, gaps in biodiversity area coverage still exist with new areas being continually added (KBA Standards and Appeals Committee 2019). Additionally, not all biodiversity areas are located in areas deemed effective for conservation but are often designated by socio-economic factors related to competing human activities (Pringle 2017; Morán-Ordóñez 2020; Rodrigues & Cazalis 2020). Key Biodiversity Areas (KBAs, BirdLife International 2020) are sites of international significance for the global persistence of biodiversity. KBAs also protect areas important for biodiversity to an extent and aim to overlap with the entire global protected area network (Donald *et al*. 2019). The KBA concept is largely based on Important Bird and Biodiversity Areas (IBAs), a template for KBAs which aim to identify and conserve sites of global importance for bird species (Donald *et al*. 2019). Indeed, the majority of terrestrial KBAs are areas designated based on birds which contain either: (1) populations of globally threatened species, (2) populations and communities of range- or biome-restricted species, or (3) substantial congregations of specific avian taxa.

Information on where to establish new KBAs can be used to identify where current protected area networks miss key bird species, and where these gaps need filling. Gap analysis is an established method to identify discontinuities in protected or biodiversity area networks (Scott *et al*. 1993) and has been effective in setting conservation planning priorities across a range of taxa (Margules & Pressey 2000). In particular, gap analysis has identified priority conservation areas for many taxa across the highly biodiverse Neotropics (e.g., de Carvalho *et al*. 2017; Bax & Francesconi 2019; Perrig *et al*. 2020). The harpy eagle (*Harpia harpyja*) is a large raptor historically distributed throughout Neotropical lowland tropical forest from southern Mexico to northern Argentina (Miranda *et al*. 2019; Sutton *et al*. 2021). The species is currently categorised as ‘Near-Threatened’ by the IUCN due to continued habitat loss and persecution (Birdlife International 2017). Harpy eagles are now largely restricted to tropical lowland broadleaf forest but can also inhabit dry seasonal forest and fragmented habitat (Vargas González *et al*. 2006; Silva *et al*. 2013).

Despite this habitat specialization, the harpy eagle has a large range due to the extensive distribution of lowland tropical forest across the Neotropical region. However, historical, and ongoing deforestation has led to localized extinctions in parts of Middle America, and the Atlantic Forest of Brazil (Vargas González *et al*. 2006; Silva *et al*. 2013; Meller & Guadagnin 2016). Current deforestation rates across the species’ stronghold in Amazonia are also of significant concern for its future persistence (Banhos *et al*. 2016; Miranda *et al*. 2019). As an apex predator requiring large tracts of continuous tropical lowland forest for breeding and foraging (Vargas González *et al*. 2014; Miranda 2015), the harpy eagle may also act as a useful trigger species for designating new regional IBAs (BirdLife International 2020), under the assumption that triggering a regional IBA would be justification for inclusion as a KBA. Further, as a Near-Threatened species of conservation concern, it fulfils the criteria for designating new regional IBAs based on inferred habitat area (category B1a; BirdLife International 2020), with the assumption that the gap sites identified are predicted to hold significant numbers of a Near-Threatened species based on preferred habitat.

Drawing from these approaches, we develop a novel predictive spatial modelling framework, first, using Resource Selection Functions to identify preferred habitat, and second, predicting habitat suitability in geographical space using a Habitat Suitability Model. Estimating harpy eagle distribution based solely on habitat predictors at the continental scale should provide the most accurate and reliable estimate of range size due to the harpy eagle’s generally high reliance on tropical lowland forest. Specifically, we set out a baseline assessment of large-scale habitat use limiting harpy eagle distribution – a first estimate of modelled habitat suitability using a new method based on the Area of Habitat metric is then used to predict areas of highest habitat suitability for the harpy eagle. Using this information, we then generate a broad-scale gap analysis to identify priority areas of highest habitat suitability in regions with limited KBA network coverage. In short, we apply statistical modelling to systematic conservation planning to determine: **(1)** how effective the current KBA network is for covering areas of harpy eagle habitat, and **(2)** where gap areas of highest habitat suitability for the harpy eagle are located for inclusion as proposed KBAs.

## Methods

### Occurrence data

Harpy eagle occurrences were sourced from the Global Raptor Impact Network (GRIN, McClure *et al*. In press), a data information system for all raptor species. For the harpy eagle, GRIN includes occurrence data from the Global Biodiversity Information Facility (GBIF 2019), which are mostly eBird records (79 %, Sullivan *et al*. 2009), along with two additional occurrence datasets (Vargas González & Vargas 2011; Miranda *et al*. 2019). Duplicate records and those with no geo-referenced location were removed and only occurrences recorded from year 2000 onwards were included to temporally match the timeframe of the habitat covariates. A 5-km spatial filter was applied between each occurrence point, which approximately matches the resolution of the raster data (∼4.5-km) and reduces the effect of biased sampling (Kramer-Schadt *et al*. 2013). A total of 1021 geo-referenced records were compiled after data cleaning. Applying the 5-km spatial filter resulted in a filtered subset of 591 harpy eagle occurrence records for use in the calibration models.

### Habitat covariates

To predict occurrence, habitat covariates representing landcover, topography and vegetation heterogeneity were downloaded from the EarthEnv (www.earthenv.org) and ENVIREM (Title & Bemmels 2018) repositories. Six continuous covariates were used at a spatial resolution of 2.5 arc-minutes (∼4.5-km resolution; See Appendix 1). Covariates were selected *a prioiri* based on the IUCN AOH model criteria from landcover and topographic factors related empirically to harpy eagle distribution and tropical forest raptor abundance in previous studies (Table 1; Robinson 1994; Anderson 2001; Vargas González & Vargas 2011; Miranda *et al*. 2019; Vargas González *et al*. 2020; Sutton *et al*. 2021). Raster layers were cropped using a delimited polygon consisting of all known range countries (including Formosa, Jujuy, Misiones and Salta provinces in northern Argentina, and Chiapas, Oaxaca, and Tabasco states in southern Mexico).

**Table 1.** Habitat covariates used in all spatial modelling analyses for the harpy eagle, with citations for the sources of the environmental data used.

| Predictor | Source | Citation |
| --- | --- | --- |
| Cultivated (%) | EarthEnv | Tuanmu & Jetz 2014 |
| Elevation (m) | EarthEnv | Amatulli *et al.* 2018 |
| Evergreen forest (%) | EarthEnv | Tuanmu & Jetz 2014 |
| Homogeneity (0.0-1.0) | EarthEnv | Tuanmu & Jetz 2015 |
| Mixed trees (%) | EarthEnv | Tuanmu & Jetz 2014 |
| Terrain Ruggedness Index (TRI) | ENVIREM | Title & Bemmels 2018 |

### Modelling

#### Resource Selection Functions

We used presence points and six covariates to fit an RSF using logistic regression with a binomial error term and logit link function using a generalised linear model (GLM) in the R stats package (R Core Team, 2018). The RSF followed geographical range first-order selection (Johnson 1980) using design I in a use-availability sampling protocol (Manly *et al*. 2002; Thomas & Taylor 2006). Linear and quadratic terms were fitted dependent on the scaled responses from fitting both terms on an initial model. That is, only linear terms were used when the quadratic term resulted in biologically unrealistic U-shaped curves, or when a linear term was sufficient to explain the scaled response. The RSF was fitted to derive maximum likelihood estimates on model parameters significantly different from zero, with no interaction terms. Predictors were standardized with a mean of zero and standard deviation of one. Because the occurrence data correspond to a presence-only dataset, background availability (which plays the role of absences) was randomly sampled using 10,000 points suitable for regression-based modelling (Barbet-Massin *et al*. 2012), and equal weights assigned to both presence and background points. To test calibration accuracy, the explained variance from each logistic model was measured using McFadden’s adjusted *R^2^* (*R^2^_adj_*, McFadden 1974). Lastly, partial response curves based on the standardized covariates of the fitted model were plotted against the scaled responses with 95 % Confidence Intervals.

We performed an Ecological Niche Factor Analysis (ENFA, Hirzel *et al*. 2002) as a second multivariate resource selection method, similar to a Principal Component Analysis (PCA), quantifying two measures of resource selection in environmental space along two axes. The first axis metric, marginality (*M*), measures the position of the species’ ecological niche, and its departure relative to the available environment. A value of *M* >1 indicates that the niche deviates more relative to the reference environmental background and has specific habitat preferences within the available environment. The second axis metric, specialization (*S*), is an inverse measure of niche breadth (the ratio of the ecological variance in mean habitat to that observed for the focal species), with a value of *S* >1 indicating higher niche specialization (narrower niche breadth). A high specialization value indicates a high reliance on the habitat conditions (variables) that mostly account for this dimension. ENFA was performed in the R package CENFA (Rinnan 2018), weighting all cells by the number of observations (Rinnan & Lawler 2019). Covariates were rescaled thus the ENFA can be interpreted similar to a PCA with eigenvalues and loadings represented along the first axis of marginality and the following, particularly the second, orthogonal axis of specialization (Basille *et al*. 2008).

### Habitat Suitability Model

We fitted HSMs using penalized elastic-net logistic regression (Zou & Hastie 2005; Fithian & Hastie 2013), in the R packages glmnet (Friedman *et al*. 2010) and maxnet (Phillips *et al*. 2017). Elastic net logistic regression imposes a regularization penalty on the model coefficients, shrinking towards zero the coefficients of covariates that contribute the least to the model, reducing model complexity (Zou & Hastie 2005; Gastón & García-Viñas 2011; Helmstetter *et al*. 2020; See Appendix 1). The maxnet package is based on the maximum entropy algorithm, MAXENT (Phillips *et al*. 2017), equivalent to an inhomogeneous Poisson process (IPP; Fithian & Hastie 2013; Renner & Warton 2013; Renner *et al*. 2015). Within the maxnet package the complementary log-log (cloglog) transform was selected as a continuous index of habitat suitability, with 0 = low suitability and 1 = high suitability. Phillips *et al*. (2017) demonstrated the cloglog transform is equivalent to an IPP and can be interpreted as a measure of relative occurrence probability proportional to a species potential abundance.

We used a random sample of 10,000 background points as pseudo-absences recommended for regression-based modelling (Barbet-Massin *et al*. 2012) and to sufficiently sample the background calibration environment (Guevara *et al*. 2018). Optimal-model selection was based on Akaike’s Information Criterion (Akaike 1974) corrected for small sample sizes (AIC_c_; Hurvich & Tsai 1989), to determine the most parsimonious model from two key MAXENT parameters: regularization multiplier (β) and feature classes (Warren & Seifert 2011). Eighteen candidate models of varying complexity were built by comparing a range of regularization multipliers from 1 to 5 in 0.5 increments, and two feature classes (response functions: Linear, Quadratic) in all possible combinations using the ‘trainMaxNet’ function in the R package enmSdm (Smith 2019). We considered all models with a ΔAIC_c_ < 2 as having strong support (Burnham & Anderson 2004), and the model with the lowest β selected to avoid overfitting. Response curves and parameter estimates were used to measure variable performance in the optimal calibration model.

We used Continuous Boyce index (CBI; Hirzel *et al*. 2006) as a threshold-independent metric of how predictions differ from a random distribution of observed presences (Boyce *et al*. 2002). CBI is consistent with a Spearman correlation (*r_s_*) and ranges from -1 to +1. Positive values indicate predictions consistent with observed presences, values close to zero suggest no difference with a random model, and negative values indicate areas with frequent presences having low environmental suitability. Mean CBI was calculated using five-fold cross-validation on 20 % test data with a moving window for threshold-independence and 101 defined bins in the R package enmSdm (Smith 2019). The optimal model was tested against random expectations using partial Receiver Operating Characteristic ratios (pROC), which estimate model performance by giving precedence to omission errors over commission errors (Peterson *et al*. 2008). Partial ROC ratios range from 0 to 2 with 1 indicating a random model. Function parameters were set with a 10% omission error rate, and 1000 bootstrap replicates on 50% test data to determine significant (*α* = 0.05) pROC values >1.0 in the R package ENMGadgets (Barve & Barve, 2013).

### Range size and gap analysis

To calculate Area of Habitat in suitable pixels and assess the effectiveness of the KBA network, we reclassified the continuous prediction to a binary threshold prediction. All pixels equal to or greater than the median pixel value of 0.345 from the continuous model were used as a suitable threshold for conservation planning (Liu *et al*. 2005; Rodríguez-Soto *et al*. 2011; Portugal *et al*. 2019). The KBA network polygons (as of September 2020; BirdLife International 2020) were then clipped to the reclassified area, establishing those KBAs covering pixels of habitat suitability ≥ 0.345 threshold. To visualise KBA network coverage, we reclassified the continuous prediction into four discrete quantile suitability classes (No habitat: 0.0 - 0.067; Low: 0.068 - 0.344; Medium: 0.345 - 0.701; High: 0.702 - 1.000).

The clipped KBA network polygons were then overlaid onto the discrete class map identifying those pixels of medium to high habitat suitability ≥ 0.345 threshold which were within the clipped KBA network polygons. We used the threshold range size to calculate a protected area ‘representation target’, quantifying how much protected area representation is needed for a species dependent on its range size following the formulation of Rodrigues *et al*. (2004),

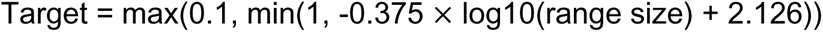

where ‘Target’ is equal to the percentage of protected target representation required for the species ‘range size’, as used in subsequent applications of the formula (Butchart *et al*. 2015; Di Marco *et al*. 2017). As can be verified by inserting different range size values, this formula yields a target of 10 % for species with a range size >250,000 km^2^ and increasing proportional representation for smaller range sizes up to a target of 100 % if range size <1000 km^2^. We used the current KBA coverage to calculate the difference between the current level of KBA coverage compared to the target level representation.

Lastly, we calculated two IUCN range metrics from our modelled AOH binary prediction. Area of Occupancy (AOO) was calculated as the number of raster pixels predicted to be occupied scaled to a 2×2 km grid following IUCN guidelines (IUCN 2018) in the R package redlistr (Lee *et al*. 2019). Extent of Occurrence (EOO) was calculated by fitting a minimum convex polygon (MCP) around the furthest boundaries of the projected habitat of the AOH polygon following IUCN guidelines (IUCN 2018). We calculated both a maximum EOO, including all the area with the MCP, and a minimum EOO, masking out the area within the MCP over the ocean. All range metric calculations were performed using an Equatorial Lambert Azimuth Equal-Area projection. General model development and geospatial analysis were performed in R (v3.5.1; R Core Team, 2018) using the dismo (Hijmans *et al*. 2017), raster (Hijmans 2017), rgdal (Bivand *et al*. 2019), rgeos (Bivand & Rundle 2019) and sp (Bivand *et al*. 2013) packages.

## Results

### Resource Selection Functions

The RSF explained 87 % of the variability in habitat use (McFadden’s *R^2^_adj_* = 0.87). Six predictors had significant terms (Table 2), with the harpy eagle most likely to be positively associated with the proportion of evergreen forest, and less likely to be associated, in declining order, with higher elevation, homogeneity, and proportion of cultivated land. Elevation had the strongest negative association, followed by homogenous vegetation, and cultivated areas (Fig. 1).

**Figure 1.**
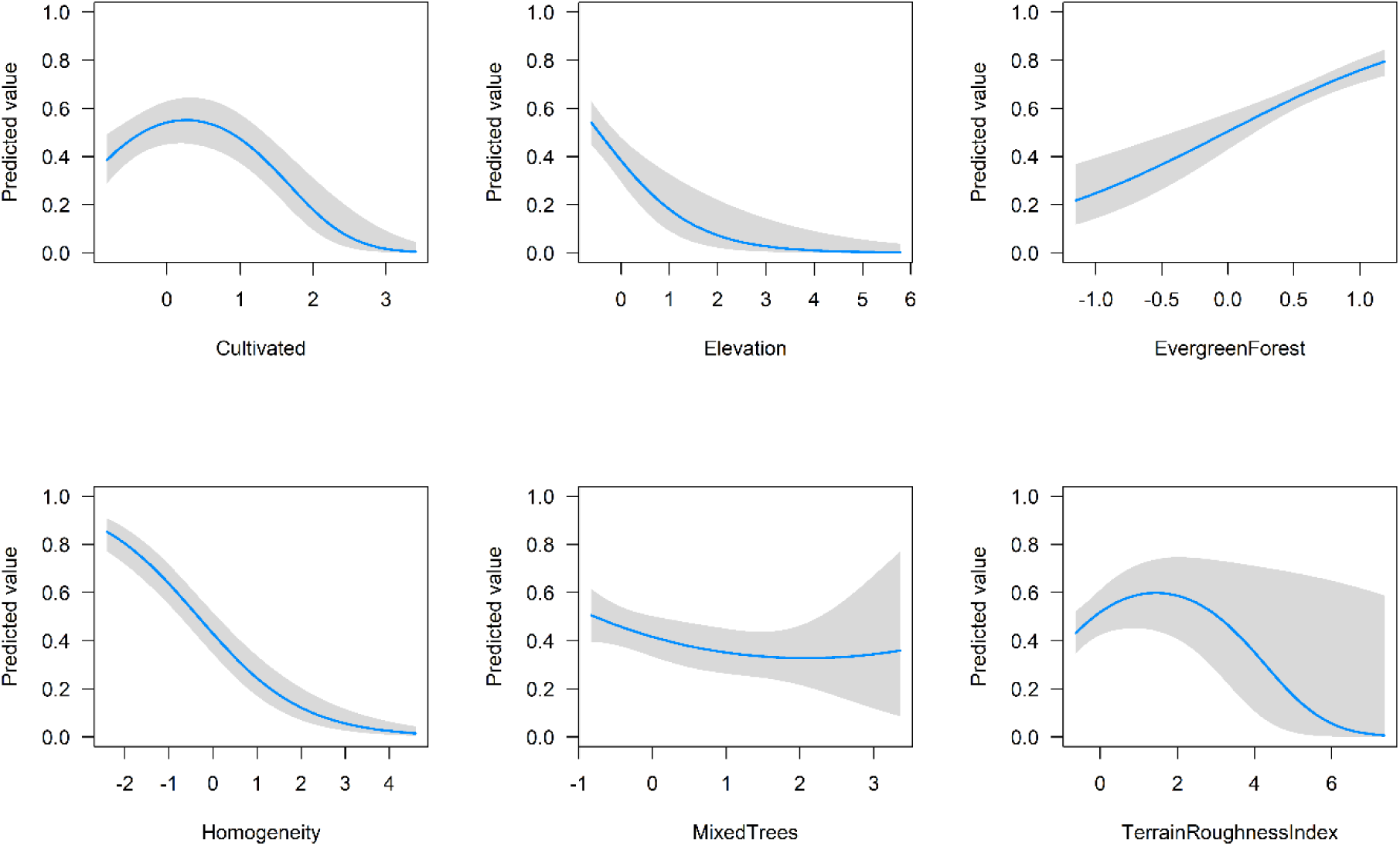
Scaled partial response curves with 95 % Confidence Intervals (grey shading) derived from maximum likelihood estimates obtained from the RSF. X-axis values are the standardized responses for each covariate (mean = 0, variance = 1).

**Table 2.** Linear and Quadratic (defined with superscript 2) terms derived from GLM maximum likelihood estimates obtained from the full model with 95 % Confidence Intervals. Covariates are ranked by the value of the regression coefficient estimates

| Predictor | Estimate | Lower CI | Higher CI | z | p |
| --- | --- | --- | --- | --- | --- |
| Intercept | -0.86 | -1.01 | -0.70 | -10.65 | 0.001 |
| Evergreen Forest | 115.33 | 89.08 | 141.58 | 8.61 | 0.001 |
| Elevation | -106.11 | -139.90 | -72.32 | -6.16 | 0.001 |
| Homogeneity | -86.84 | -101.07 | -72.61 | -11.96 | 0.001 |
| Cultivated<sup>2</sup> | -57.49 | -76.45 | -38.53 | -5.94 | 0.001 |
| Cultivated | -33.20 | -58.73 | -7.67 | -2.55 | 0.042 |
| Mixed Trees | -28.84 | -50.67 | -7.01 | -2.59 | 0.062 |
| Terrain Roughness Index<sup>2</sup> | -27.79 | -51.38 | -4.21 | -2.31 | 0.017 |
| Mixed Trees<sup>2</sup> | 7.62 | -8.74 | 23.98 | 0.91 | 0.478 |
| Terrain Roughness Index | 0.26 | -25.27 | 25.79 | 0.02 | 0.987 |

The ENFA habitat space deviated from the average background habitat available (Fig. 2), with the marginality factor higher than the average background habitat (*M =* 1.026) accounting for 11.37% of the total variance (Table S2). The harpy eagle was restricted to specific habitat relative to the background habitat with specialized habitat requirements (*S* = 1.886). Four significant ENFA factors explained 86.19 % of total variance, with the first specialization axis (Spec1) explaining 43.78 % (Table S2). Evergreen forest had the highest coefficient on the marginality axis (0.69), and elevation the highest on the specialization axis (0.89).

**Figure 2.**
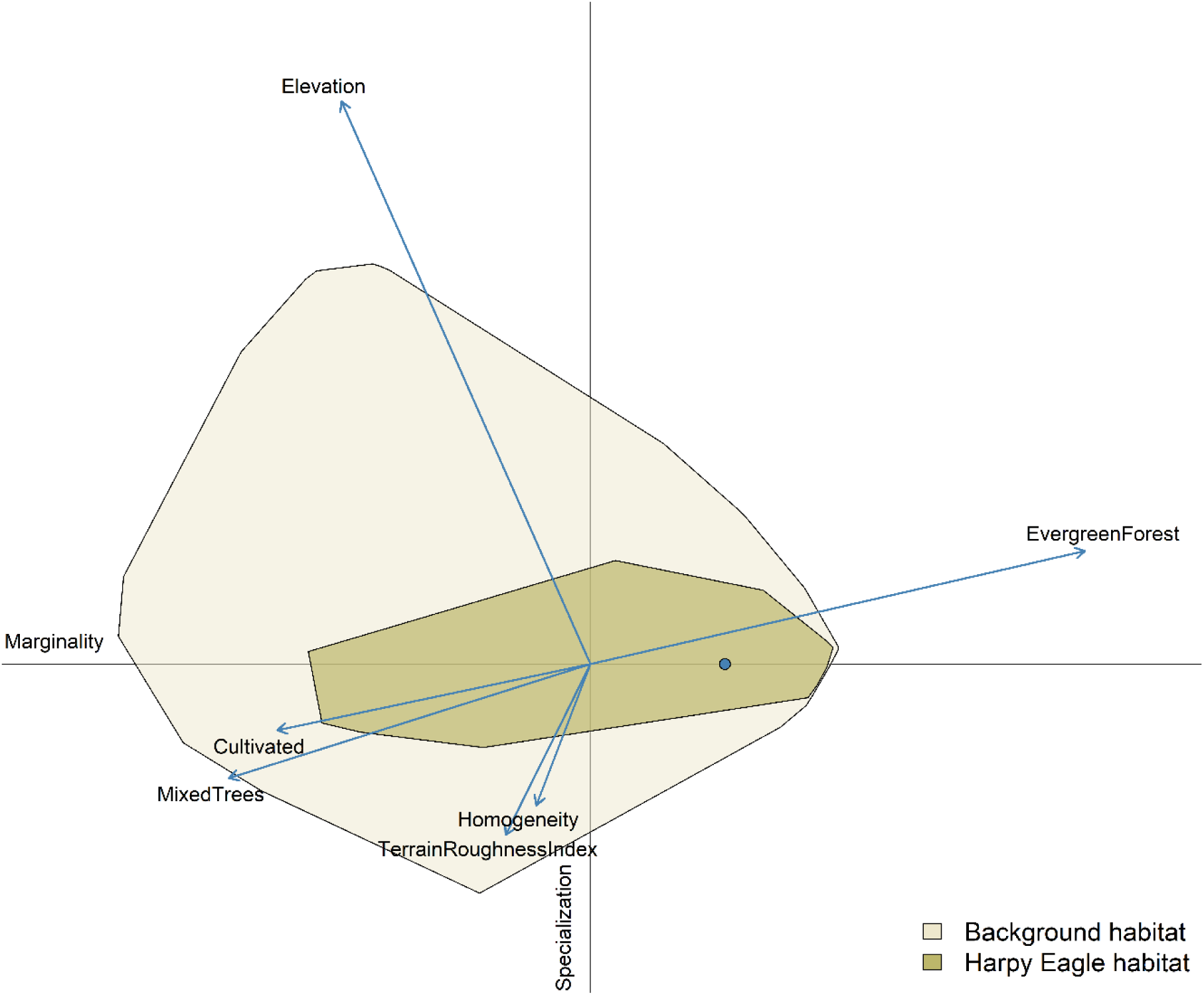
Ecological Niche Factor Analysis (ENFA) for suitable harpy eagle habitat space (khaki) within the available background environment (beige) shown across the marginality (x) and specialization (y) axes. Blue point indicates niche position (median marginality) relative to the average background environment (the plot origin). Arrow length indicates the magnitude with which each covariate accounts for the variance on each of the two axes. For example, Evergreen Forest and Elevation are the two most important covariates on the marginality and specialization axes respectively. Both have longer arrow lengths representing a higher percentage of the variance accounted for in the eigenvalues relative to the remaining covariates.

### Habitat Suitability Model

Six candidate models had an ΔAIC_c_ ≤ 2, and the model with the lowest regularization multiplier (β) was selected (Model 6 in Table S3). The best-fit HSM (ΔAIC_c_ = 1.19) had linear and quadratic terms and β = 2.5 as model parameters, with high calibration accuracy (mean CBI = 0.960), and was robust against random expectations (pROC = 1.431, SD± 0.055, range: 1.244 – 1.594). Harpy eagles were most positively associated with evergreen forest and negatively associated with habitat homogeneity (Table 3). The largest continuous area of habitat extended across Amazonia and the Guiana Shield (Fig. 3). A habitat corridor was identified through Central America along the Caribbean coast, extending south into the Chocó-Darién ecoregion along the Pacific coast of Colombia. Little habitat was predicted across the largely deforested Atlantic Forest region in Brazil. From the HSM response curves, evergreen forest had peak suitability at 70-75 % forest cover, with highest suitability for topographic areas of both low elevation and terrain ruggedness (Fig. 4). Habitat suitability was highest in areas of low homogeneity < 0.2 (i.e., highly heterogenous vegetation), areas with < 10 % human cultivated landcover, and zero or low percentage of mixed trees (i.e., mosaic forest and shrub/grassland).

**Figure 3.**
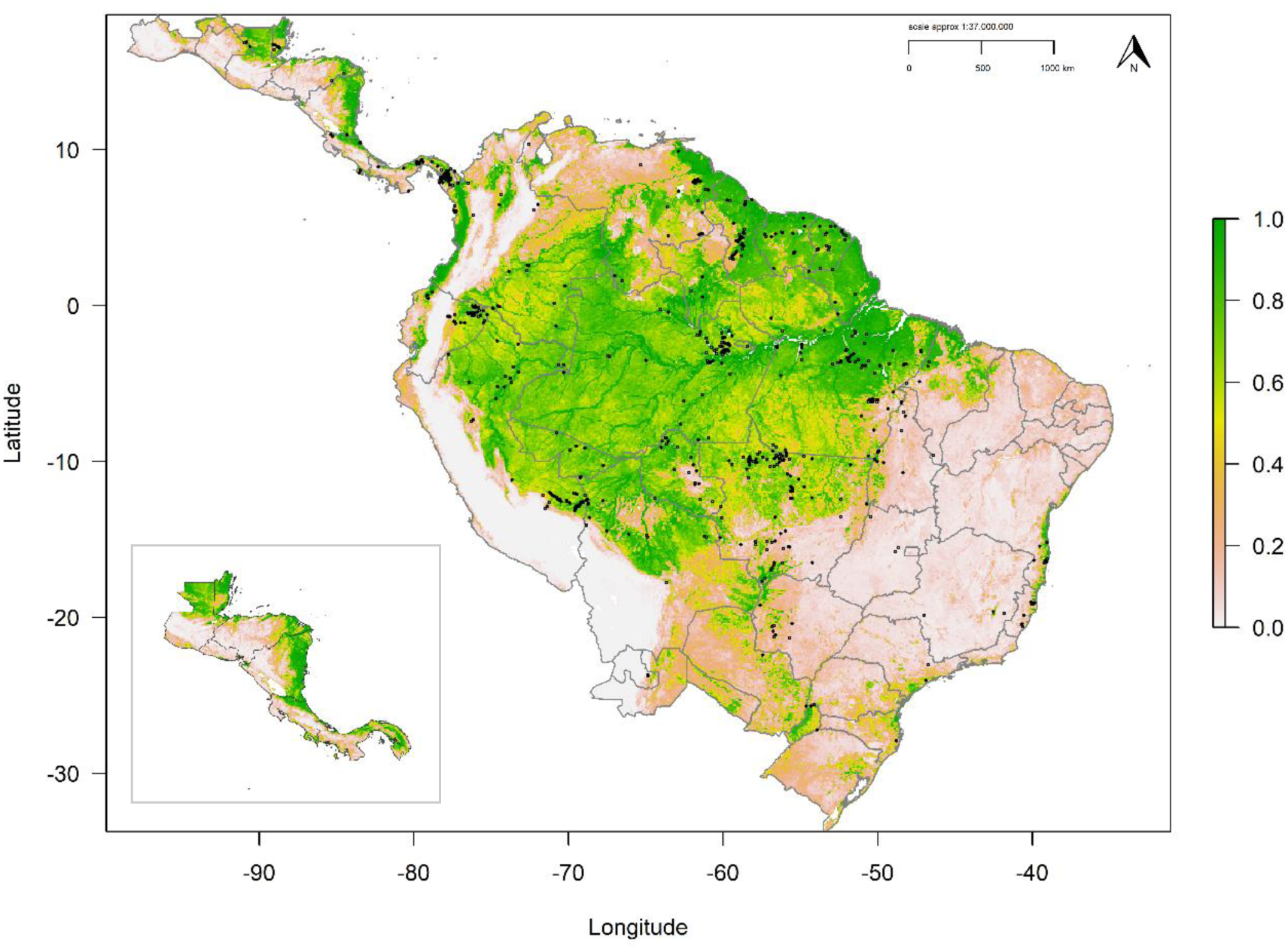
Habitat Suitability Model for the harpy eagle. Map denotes cloglog prediction with darker green areas (values closer to 1) having highest suitability and expected abundance. Grey borders define national boundaries and internal state boundaries for Argentina, Brazil, and Mexico. Black points define harpy eagle occurrences. Inset map shows cropped model prediction for Central America without harpy eagle occurrences for clarity.

**Figure 4.**
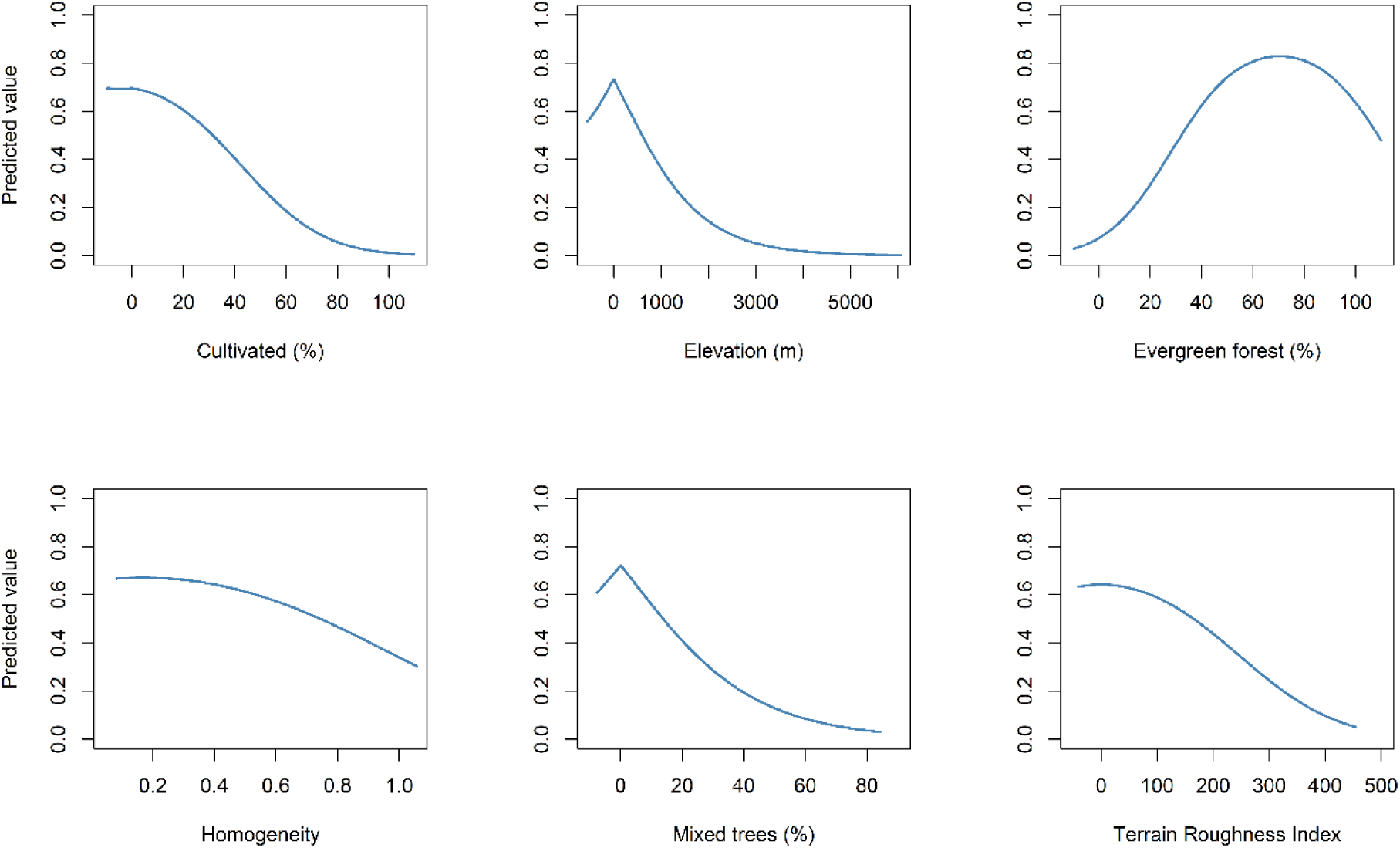
Covariate response curves from the Habitat Suitability Model for the harpy eagle.

**Table 3.**
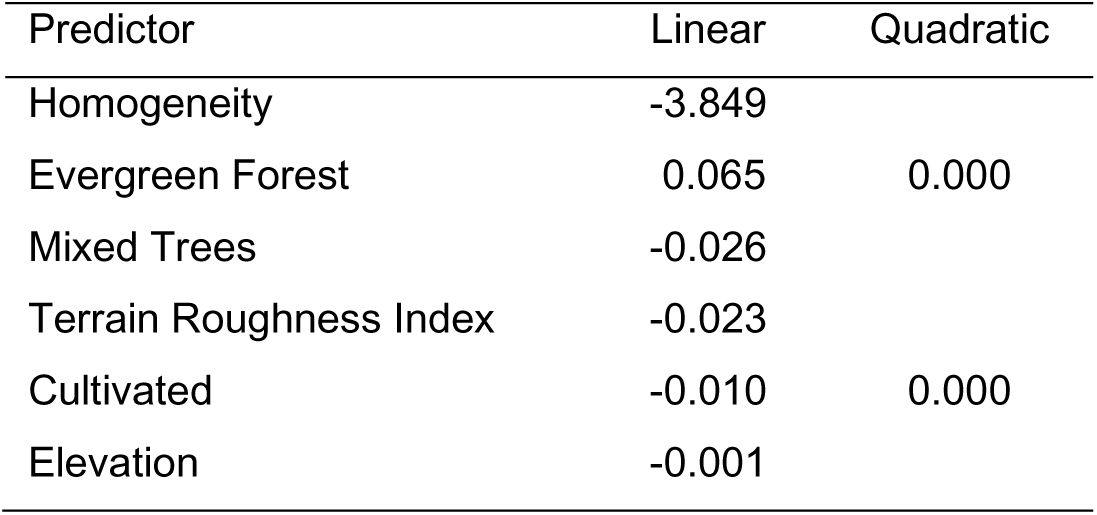
Parameter estimates derived from the HSM penalized elastic net regression beta coefficients.

### Range size and gap analysis

The reclassified binary model (median threshold = 0.345) calculated an Area of Habitat equalling 7,479,752 km^2^ (Fig. S1). The current KBA network covered 18.1 % (1,352,879 km^2^) of this habitat area in the medium to high discrete quantile classes (Fig. 5), 8.1 % greater than the target representation (10 %). Four major gaps (Fig. 5, blue circles/ellipses) for high threshold habitat without extensive KBA coverage were identified in: (**1**) the Chocó-Darién ecoregion in western Colombia (Fig. 6), (**2**) the Magdalena-Urabá moist forests of northern Colombia (Fig. 6), (**3**) north-east Amazonas state in Brazil, and (**4**) north and west Guyana. From our AOH model, maximum EOO was 18,130,602 km^2^ and minimum EOO 14,738,408 km^2^, with an AOO of 708,697 occupied cells.

**Figure 5.**
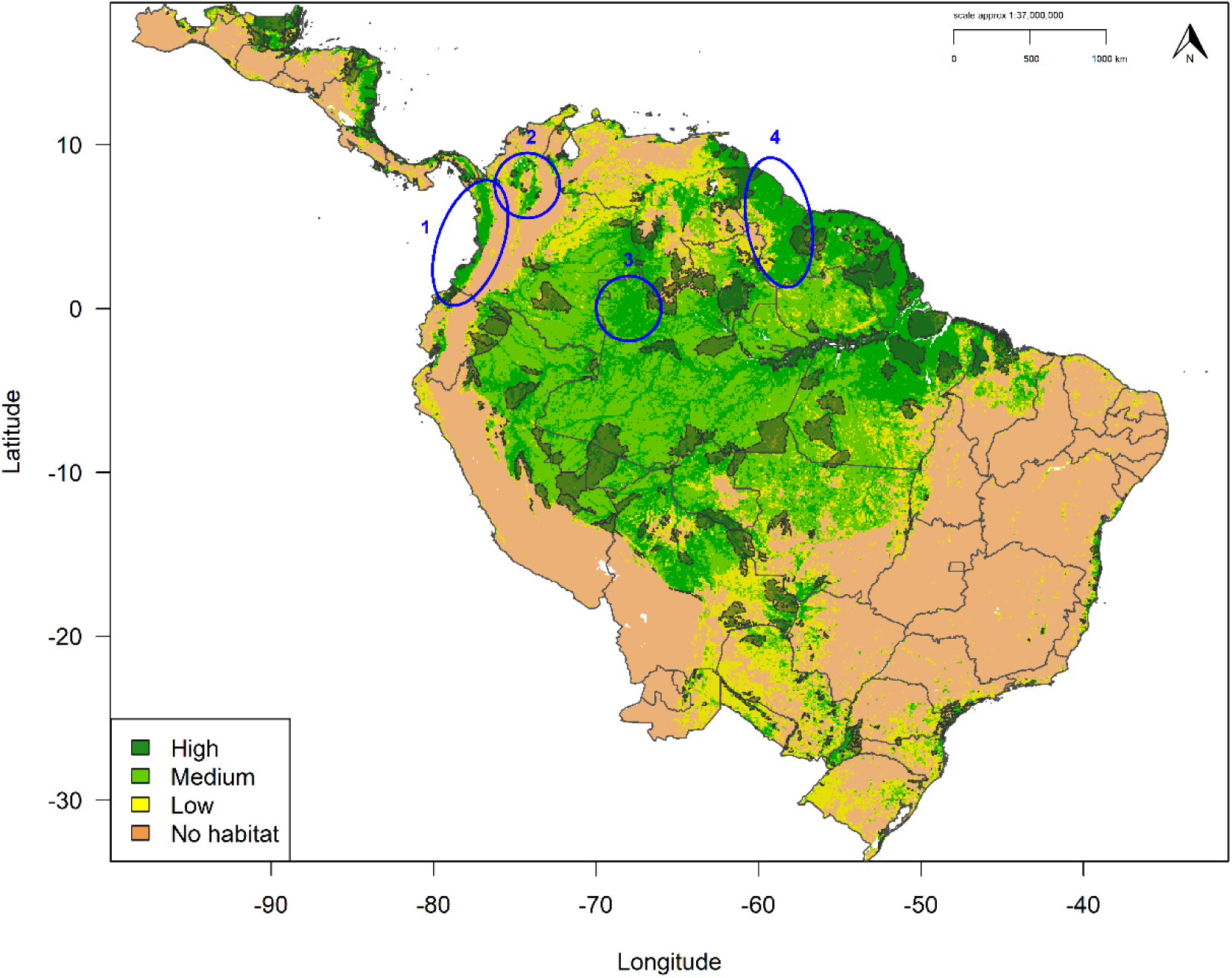
Key Biodiversity Area (KBA) network gap analysis for harpy eagle habitat. Map denotes cloglog prediction reclassifed into four discrete quantile threshold classes (brown = no habitat; yellow = low, pale green = medium; dark green = high). Black bordered polygons denote current KBA network. Blue ellipses identify priority KBA network coverage gaps: (**1**) Chocó-Darién region in Colombia, (**2**) Magdalena-Urabá moist forests in northern Colombia, (**3**) north-east Amazonas state in Brazil, (**4**) north and west Guyana. Grey borders define national boundaries and internal province and state boundaries for Argentina, Brazil, and Mexico.

**Figure 6.**
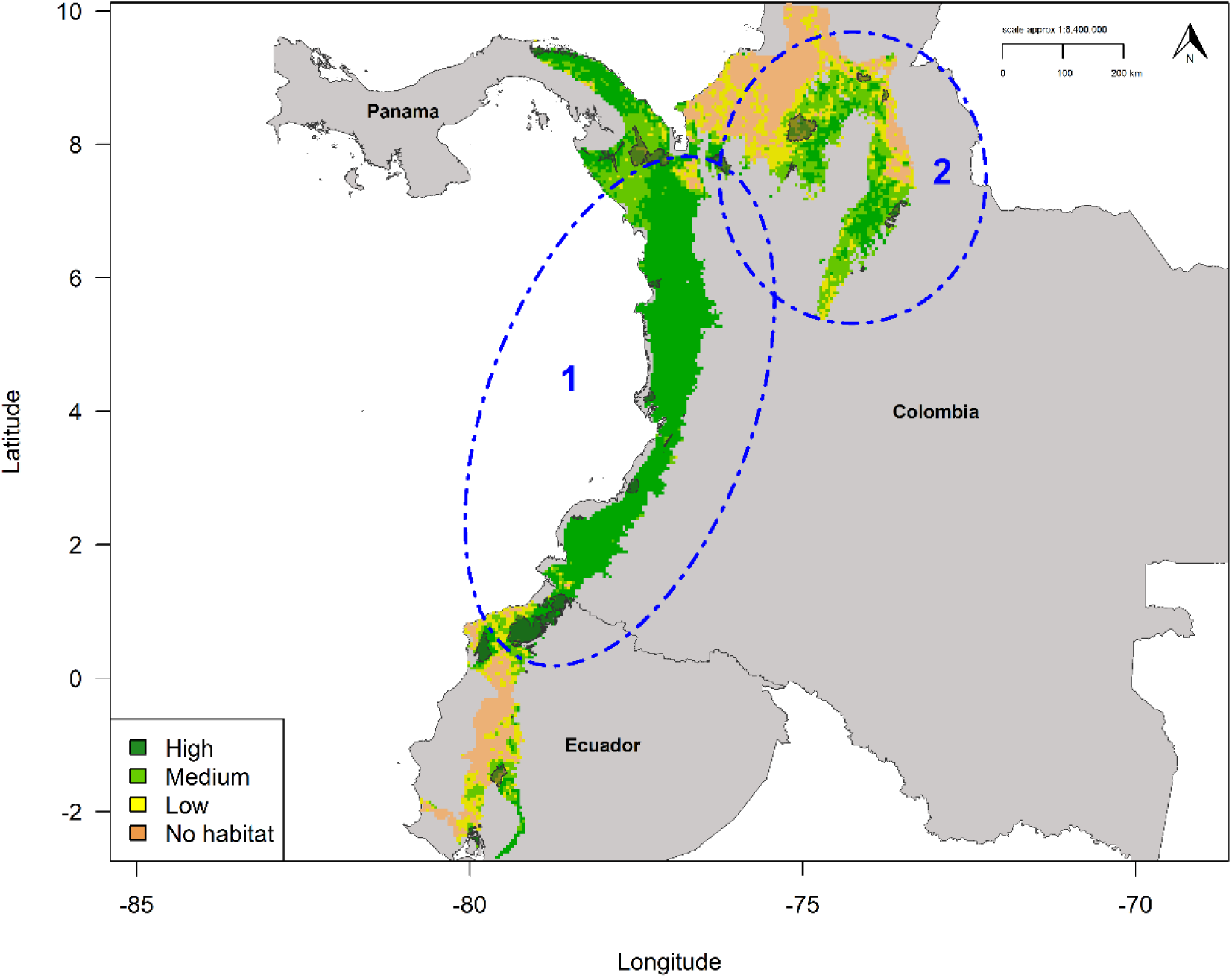
Key Biodiversity Area (KBA) network gap analysis for harpy eagle habitat projected into the Chocó-Darién ecoregion. Map denotes cloglog prediction reclassifed into four discrete quantile threshold classes (brown = no habitat; yellow = low, pale green = medium; dark green = high). Black bordered transparent polygons denote current KBA network. Hashed blue ellipses identify priority KBA network coverage gaps: (**1**) Chocó-Darién region in Colombia, Ecuador, and Panama, (**2**) Magdalena-Urabá moist forests in northern Colombia.

## Discussion

Our results indicate that harpy eagle populations are more likely to be associated with dense (70-75%) evergreen forest cover, low elevation, and high vegetation heterogeneity across their range. Conversely, harpy eagles seem to avoid extensive areas of cultivated land, mosaic forest, and high terrain complexity. Using the AOH parameters as the basis for the habitat model predicted a large area of habitat across the pan-Amazonia region, and a habitat corridor extending from the Pacific coast of Colombia, north along the Caribbean coast of Central America. Almost no habitat was predicted across the Atlantic Forest region, which is now severely degraded. The current KBA network coverage exceeded the target representation (10 %), covering ∼18 % of medium to high harpy eagle habitat. Considering the large range of the harpy eagle, the current KBA extent is encouraging but misses key areas of potentially important habitat. Four areas of high habitat suitability were identified as gaps in the KBA network for north and west Colombia, western Guyana, and north-west Brazil. We recommend establishing new KBAs in these four areas, further strengthening the current KBA network across the region.

### Habitat use

Broad and fine scale species-habitat assessments often result in different variables emerging as important, potentially leading to contrasting recommendations for conservation (Gregory & Baillie 1998). However, our results show general similarities to habitat models from previous studies at both broad and fine scales. The HSM was consistent with predicted harpy eagle habitat from an earlier broad-scale HSM (Miranda *et al*. 2019). This was expected because both HSMs used measures of forest cover as landcover predictors but with differing modelling methodologies. This reinforces the consistency in HSM outputs for the harpy eagle from a range of algorithms and gives confidence in HSM predictions that have been criticised for lacking ecological realism (Fourcade *et al*. 2017). Building on the Miranda *et al*. (2019) model, the HSM here also predicted a distinct corridor of habitat extending from the Chocó-Darién ecoregion of west Colombia north through Central America along the Caribbean coast (Fig. 6). This suggests that including a habitat heterogeneity covariate, along with topographic and landcover predictors, was able to identify key areas of habitat undetectable from other texture measures used in that study.

Habitat heterogeneity is a key landscape characteristic, important for determining general biodiversity patterns (Stein *et al*. 2014), including for lowland tropical forest raptors (Jullien & Thiollay 1996; Anderson 2001). Areas of higher heterogeneity provide more environmental niche space, promoting greater species coexistence and thus increased species diversity (Tews *et al*. 2004). For the harpy eagle, areas of higher habitat heterogeneity may be preferred over more homogenous areas, as they contain a greater density and diversity of prey species (Miranda 2018). Further, a diverse forest canopy structure may also facilitate aerial attacks on canopy prey, by providing more hunting perches (Vargas González *et al*. 2014). Moreover, the HSM confirmed the restricted elevational distribution for the harpy eagle, consistent with a landscape-level HSM (Vargas González *et al*. 2020). This may be similarly linked to the harpy eagles’ preference for nesting in large, canopy-emergent trees, and the abundance of its main prey of arboreal mammals, both of which occur in greater abundance at lower elevations (Miranda 2015; Miranda *et al*. 2020).

Harpy eagles are dependent on large tracts of lowland tropical forest for breeding and foraging (Vargas González *et al*. 2014; Miranda *et al*. 2019). Indeed, breeding success was higher in areas with > 70 % forest cover in northern Mato Grosso, Brazil (Miranda *et al*. 2021), consistent with the range-wide response to evergreen forest cover here. Perhaps as important, strong negative associations were identified with >10 % cultivated landcover and mosaic forest, showing that harpy eagles avoid areas of high human impact. This implies that, as deforestation increases across the species’ range, the harpy eagle may struggle to adapt to large areas of human disturbance and heavily fragmented landscapes (Miranda *et al*. 2021).

### Area of Habitat

Our new method of calculating the Area of Habitat metric refines previous range size estimates (Birdlife International 2017; Sutton *et al*. 2021) and provides a new basis for an updated range map for the harpy eagle. There was 32.4 % less area in our modelled AOH range polygon (7,479,752 km^2^), than in the current IUCN range map polygon (11,064,113 km^2^; Fig. S1). Therefore, we recommend this new range size estimate be incorporated into future IUCN assessments for the species. Our modelled AOH polygon also had 24 % less area compared to an HSM using solely climatic and topographic predictors (9,844,399 km^2^; Sutton *et al*. 2021). If we assume that the HSM from Sutton *et al*. based on climate and terrain is representative of the harpy eagle pre-industrial range (in the absence of satellite-derived landcover not available for pre-industrial times), then the species’ habitat range has shrunk by nearly a quarter during the industrial period to the present. One limitation of the analyses was the timeframe of the remote-sensing data used for the covariates. Both the landcover and vegetation covariates are a consensus product collected between the years 2000-2005, with land use having changed in parts of Neotropics since then (Powers & Jetz 2019). Therefore, the area of habitat prediction should be viewed as a baseline assessment, knowing that landcover can change rapidly. However, processing large areas of current remote-sensed landcover data at continental-scales can be problematic due to the high computing power required. Even so, the EarthEnv habitat variables are recommended as a readily available dataset to use for first estimates of modelled AOH at large scales (Tuanmu & Jetz 2014, 2015).

Current and predicted future habitat loss may lead inevitability to declines in populations of some species, increasing their extinction risk (Powers & Jetz 2019). Continued habitat loss and fragmentation is likely to have a negative impact on the future persistence for many birds across the highly biodiverse Neotropics (Bird *et al*. 2011). The harpy eagle is a good example, despite its large range precluding high extinction risk (Gaston & Fuller 2009). Continued habitat loss and fragmentation through agricultural development and logging across its geographic range (Vargas González *et al*. 2006; Miranda *et al*. 2020) should raise the alarm about the species’ future (Krüger & Radford 2008; Miranda *et al*. 2019). This is demonstrated by the few harpy eagle breeding and sighting records in the largely deforested Atlantic Forest (Meller & Guadagnin 2016; Suscke *et al*. 2017), and parts of Meso- and Central America (Vargas González *et al*. 2006), which is reflected in the results from the HSM. Our results should therefore serve as a fore-warning of what could happen across parts of the core habitat area in Amazonia where deforestation has steadily increased since 2000 (Hansen *et al*. 2008).

As a baseline assessment, our HSM should be viewed as a *maximum extent of habitat*, knowing that deforestation is an ongoing process across the pan-Amazonia region (Bird *et al*. 2011; Hansen *et al*. 2020). Approximately 18 % of tropical forest in Amazonia had been cleared by 2011 (Bird *et al*. 2011), with predictions of up to 40 % of forest cover lost by 2050 (Soares-Filho *et al*. 2006). Recently, those tropical forests of highest structural integrity most associated with preferred harpy eagle habitat (tall, closed canopy forest and low human pressure; Vargas González *et al*. 2014; Miranda *et al*. 2020) were identified as largely limited to the Amazon basin (Hansen *et al*. 2020). These forests generally remain intact due to their remoteness (Soares-Filho *et al*. 2006), but with the majority having no formal protection. Strengthening biodiversity and protected area networks should be given high priority in policy decisions (Butchart *et al*. 2015), along with effective biodiversity area-based conservation outside of, but concurrent with, formally protected areas (Pringle 2017; Maxwell *et al*. 2020).

### Gap analysis

Although the current coverage of the KBA network within our modelled AOH range (∼18 %) exceeded the representative target set here (10 %), it is substantially lower than the proportion of IBA network coverage for threatened bird species overall in Amazonia (54.9 %, Bird *et al*. 2011). Of the four key gaps identified here only gap 3 in north-west Amazonas state in Brazil has any form of current protection as an area of indigenous land (UNEP-WCWC 2020). The three remaining gap areas have little formal protection or KBA coverage, despite both the Chocó-Darién ecoregion (gap 1) and Guyana (gap 4) having extensive harpy eagle habitat. In the case of north and west Guyana it is likely that most habitat is ‘passively’ protected due to the inaccessibility of the region. However, solely relying on remoteness may be short-sighted and extending the current KBAs east and west of Guyana to cover a larger portion of the Guiana Shield is recommended. To this aim, given that on average ∼49 % of the area of each KBA/IBA globally has formal protection (Waliczky *et al*. 2019), intersecting KBA coverage with nationally protected areas across the harpy eagle range would be a useful next step in protected area assessment for the species (Butchart *et al*. 2012).

The Chocó-Darién ecoregion is one of 25 global biodiversity hotspots prioritized for conservation (Myers *et al*. 2000). Based on satellite remote-sensing, deforestation for agricultural expansion has steadily increased in the region over the past two decades (Fagua *et al*. 2019; Fagua & Ramsey 2019). Approximately 42 % of forest remains intact, making this an area of high importance for protection not only for the harpy eagle but for all the associated fauna, flora, and crucial ecological processes. Establishing and reinforcing the current KBA network throughout the Chocó-Darién ecoregion could be important for habitat continuity essential to dispersing harpy eagles (Urios *et al*. 2017) between Central and South America. The Darién region of Panama has a high density of breeding harpy eagles and is considered the current stronghold of the species in Central America (Vargas González & Vargas 2011). Designating new KBAs in the Chocó-Darién ecoregion corridor could thus sustain habitat for fragmented harpy eagle populations maintaining genetic diversity and thus potential adaptation to environmental change (Lerner *et al*. 2009; Banhos *et al*. 2016; Maxwell *et al*. 2020). Indeed, genetic diversity decreased in fragmented harpy eagle populations inhabiting deforested regions of the southern Amazon and Atlantic Forest of Brazil (Banhos *et al*. 2016), reinforcing the need to protect and link habitat patches throughout its whole distribution.

Habitat loss is a principal threat to the long-term survival of the harpy eagle and protecting large areas of tropical forest habitat for the species should be a high priority (Banhos *et al*. 2016). Continued deforestation resulting in habitat loss and fragmentation across the harpy eagle range should raise the alarm about the species’ future conservation status. Using targeted forest protection through responsible community land use and broad-scale conservation planning is needed to reduce current deforestation rates (Kramer *et al*. 1997; Bird *et al*. 2011; Butchart *et al*. 2015). Whilst the current KBA network coverage for the harpy eagle exceeds the representation target, our models identified gaps in the KBA network that ought to be prioritised for enlarging the KBA network estate. As demonstrated here, our new method of calculating modelled area of habitat estimates based on HSMs are a useful tool for large-scale conservation planning and can be readily applied to many taxa.

## Acknowledgments

We thank all individuals and organisations who contributed occurrence data to the Global Raptor Impact Network (GRIN) information system. LJS thanks The Peregrine Fund for providing financial assistance for his doctoral studentship. EBPM work is supported by Rufford Foundation, ONF Brasil, Rainforest Biodiversity Group, Idea Wild, Explorer’s Club Exploration Fund, Cleveland Metroparks Zoo, and SouthWild.com. We thank the M.J. Murdock Charitable Trust for funding and the Information Technology staff and Research Library interns at The Peregrine Fund for support.

## Data Accessibility Statement

Upon acceptance the data that support the findings of this study will be made openly available on the data repository *figshare*

## Appendices

## Appendix 1 Supplementary Methods

### Habitat covariates

Elevation and Terrain Roughness Index (TRI) are both key topographic variables driving harpy eagle distribution (Vargas González & Vargas 2011; Vargas González *et al*. 2020; Sutton *et al*. 2021). Elevation was derived from a digital elevation model (DEM) product from the 250m Global Multi-Resolution Terrain Elevation Data 2010 (GMTED2010, Danielson & Gesch 2011). TRI was derived from the 30 arc-sec resolution Shuttle Radar Topographic Mission (SRTM30, Becker *et al*. 2009). Homogeneity is a similarity measure for Enhanced Vegetation Index (EVI) between adjacent pixels; sourced from the Moderate Resolution Imaging Spectroradiometer (MODIS, https://modis.gsfc.nasa.gov/). Homogeneity varies between zero (zero similarity = maximum heterogeneity) and one (complete similarity).

The three measures of percentage landcover (Evergreen Forest, Mixed Trees, Cultivated) are consensus products integrating GlobCover (v2.2), MODIS land-cover product (v051), GLC2000 (v1.1) and DISCover (v2) at 30 arc-sec (∼1km) spatial resolution. Mixed trees represents a mosaic landcover of forest, shrubland and grassland. All landcover layers were resampled to a spatial resolution of 2.5 arc-minutes using bilinear interpolation. Full details on methodology and image processing can be found in Tuanmu & Jetz (2014) for the landcover layers, and Tuanmu & Jetz (2015) for the habitat heterogeneity texture measures. All selected covariates showed low collinearity and thus all six were included as predictors in model calibration (Variance Inflation Factor (VIF) < 5; Table S1).

### Habitat Suitability Model

In its original implementation MAXENT imposed a ‘lasso’ (least absolute shrinkage and selection operator) regularization penalty, where only the most significant covariates are retained, with uninformative covariates set at zero. Instead, the maxnet package uses an elastic net to perform automatic covariate selection (lasso) and continuous shrinkage (ridge regression) simultaneously (Zou & Hastie 2005; Phillips *et al*. 2017), evaluating the contribution of all covariates and shrinking low-contribution coefficients towards zero. Elastic net regularization improves predictive accuracy compared to the lasso, in both simulated and real data examples (Zou & Hastie 2005) and may be viewed as a generalization of the lasso.

## Appendix 2 Supplementary Tables

**Table S1.** Multi-collinearity test using stepwise elimination Variance Inflation Factor (VIF) analysis. Variables with VIF < 5 have low correlation with other variables, and thus are suitable for inclusion in calibration models when further evaluated for ecological relevance.

| Predictor | VIF |
| --- | --- |
| Homogeneity | 1.65 |
| Terrain Ruggedness Index | 1.76 |
| Elevation | 2.41 |
| Mixed trees | 2.54 |
| Cultivated | 2.62 |
| Evergreen forest | 4.64 |

**Table S2.** Coefficient values (eigenvectors) for the marginality factor and the first three specialization factors from the ENFA analysis, ordered from highest to lowest on the marginality factor, and percentage of variance accounted for by each of these factors (Marg. = marginality; Spec = Specialization)

| ENFA axis | Marg. | Spec1 | Spec2 | Spec3 |
| --- | --- | --- | --- | --- |
| Variance explained (%) | 11.37 | 43.78 | 18.01 | 13.03 |
| Predictor |  |  |  |  |
| Evergreen forest | 0.69 | 0.18 | -0.62 | 0.56 |
| Mixed trees | -0.50 | -0.18 | -0.14 | 0.72 |
| Cultivated | -0.43 | -0.11 | -0.65 | -0.13 |
| Elevation | -0.35 | 0.89 | -0.32 | 0.31 |
| Terrain Roughness Index | -0.12 | -0.27 | 0.19 | -0.17 |
| Homogeneity | -0.08 | -0.22 | 0.18 | -0.13 |

**Table S3.** Model selection metrics for all six candidate models with ΔAIC_c_ < 2. RM = regularization multiplier (β), FC = feature classes, LQ = Linear, Quadratic.

| Model | RM | FC | AIC<sub>c</sub> | $\Delta AIC_c$ |
| --- | --- | --- | --- | --- |
| 1 | 4.0 | LQ | 7574.316 | 0.000 |
| 2 | 3.5 | LQ | 7574.389 | 0.070 |
| 3 | 4.5 | LQ | 7574.561 | 0.245 |
| 4 | 3.0 | LQ | 7574.785 | 0.470 |
| 5 | 5.0 | LQ | 7575.125 | 0.809 |
| 6 | 2.5 | LQ | 7575.509 | 1.193 |

## Appendix 3 Supplementary Figures

**Figure S1.**
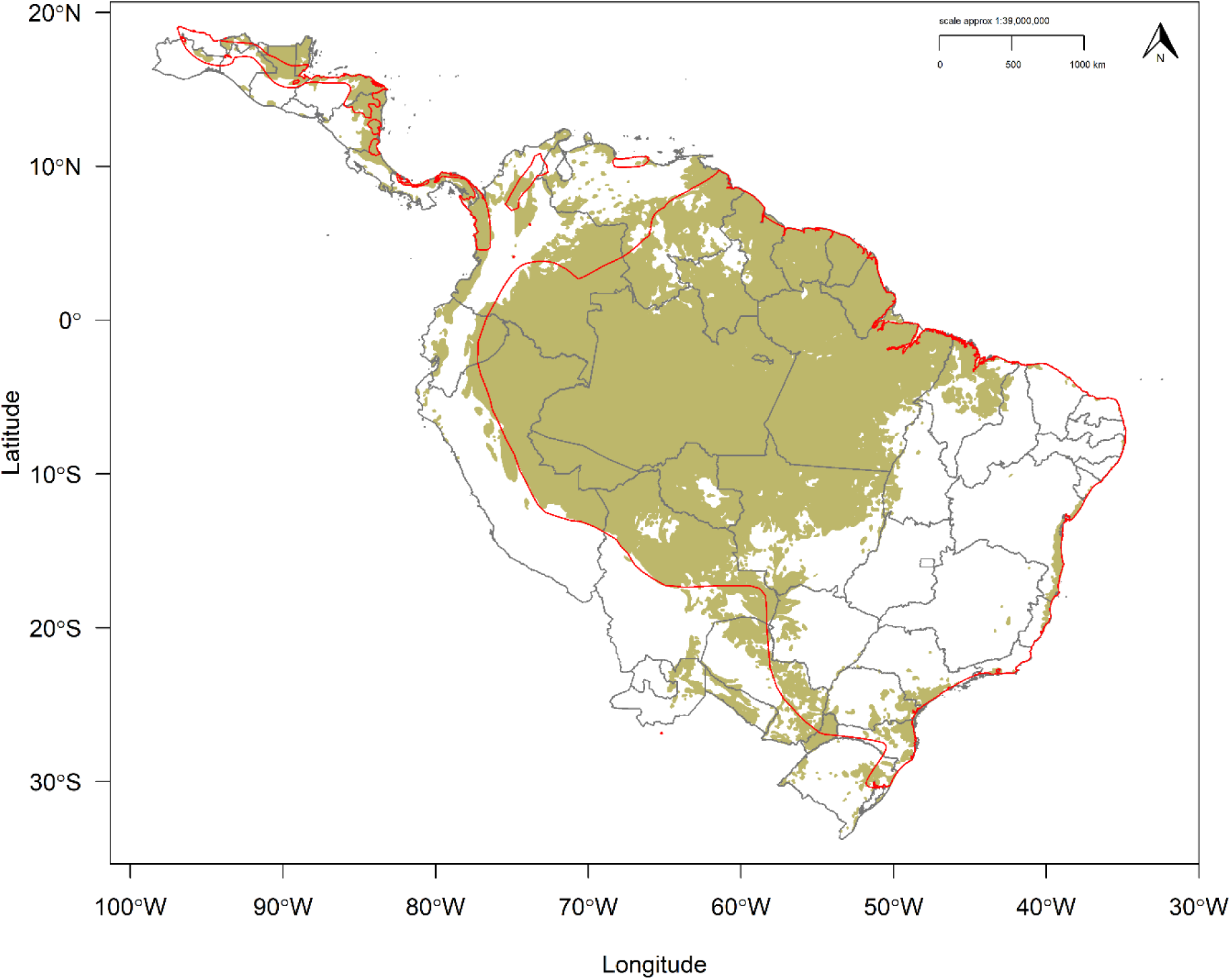
Reclassified binary Habitat Suitability Model (threshold = 0.345) for the harpy eagle. Dark khaki area is habitat above the 0.345 threshold, white areas below the threshold. Red polygons define current IUCN range map for the harpy eagle as a comparison to the HSM prediction. Grey borders define national boundaries and internal state boundaries for Argentina, Brazil, and Mexico.

